# Nascent flagellar basal bodies are immobilized by rod assembly in *Bacillus subtilis*

**DOI:** 10.1101/2024.08.02.606393

**Authors:** Caroline M. Dunn, Daniel Foust, Yongqiang Gao, Julie S. Biteen, Sidney L. Shaw, Daniel B. Kearns

## Abstract

Flagella are complex, trans-envelope nanomachines that localize to species-specific cellular addresses. Here we study the localization dynamics of the earliest stage of basal body formation in *Bacillus subtilis* using a fluorescent fusion to the C-ring protein FliM. We find that *B. subtilis* basal bodies do not exhibit dynamic subunit exchange and are largely stationary at steady state, consistent with flagellar assembly through the peptidoglycan. Rare basal bodies were observed to be mobile however, and the frequency of basal body mobility is elevated both early in basal body assembly and when the rod is mutated. Thus, basal body mobility is a precursor to patterning and we propose that rod polymerization probes the peptidoglycan superstructure for pores of sufficient diameter that permit rod completion. Furthermore, mutation of the rod also disrupts basal body patterning in a way that phenocopies mutation of the cytoplasmic flagellar patterning protein FlhF. We infer that conformational changes in the basal body exchange information between rod synthesis and the cytoplasmic patterning proteins to restrict assembly at certain pores established by a grid-like pattern pre-existent in the peptidoglycan itself.

**IMPORTANCE:** Bacteria insert flagella in a species-specific pattern on the cell body, but how patterns are achieved is poorly understood. In bacteria with a single polar flagellum, a marker protein localizes to the cell pole and nucleates the assembly of the flagellum at that site. *Bacillus subtilis* assembles ∼15 flagella over the length of the cell body in a grid-like pattern and lacks all proteins associated with targeted assembly in polarly flagellated bacteria. Here we show that *B. subtilis* basal bodies are mobile soon after assembly and become immobilized when the flagellar rod transits the peptidoglycan wall. Moreover, defects in the flagellar rod lead to an asymmetric distribution of flagella with respect to the midcell. We conclude that the patterning of flagella is different in *B. subtilis*, and we infer that the *B. subtilis* rod probes the peptidoglycan for holes that can accommodate the machine.

## INTRODUCTION

Subcellular localization is an important mechanism to control the amount and function of protein complexes. In bacteria, the subcellular localization of proteins became widely recognized when TEM immunogold-staining showed that the cell division protein FtsZ localized to the division plane and methyl accepting chemotaxis proteins localized as clusters near cell poles (1–3). An example of subcellular localization in bacteria that predates the localization of FtsZ by many decades however, is the patterning of the bacterial flagellum. Bacteria have long been known to assemble a species-specific number of flagella ranging from one to many per cell, localized to poles or along the length of a rod shaped body (peritrichous), and phylogenetic conservation suggests that both traits are selectable (4–6). Precisely how bacteria arrive at their species-specific number and location of flagella is poorly understood, and the two phenomena may be related (7,8). Flagella are complex, sequentially-assembled trans-envelope nanomachines (9–11) and patterning is likely determined early in synthesis.

Flagellar synthesis begins with the assembly of a transmembrane basal body composed of a type III secretion apparatus surrounded by the flagellar baseplate protein FliF (12–17).

Next, a cytoplasmic ring of proteins (the C-ring) is assembled onto the membrane complex that forms a gear-like rotor and directional control system (18–21). Once the C-ring is assembled and basal body formation is complete, the flagellar type III secretion system (FT3SS) becomes active and secretes subunits that polymerize atop the basal body to form the axle-like rod (12,13,19,22,23). The rod extends away from the cell and transits the peptidoglycan, as well as the outer membrane, if present (19,24). The FT3SS also exports subunits of the universal joint-like hook, and later, subunits of the long helical filament, which when rotated, serves as a propeller (25,26). Complex regulatory networks ensure that the genes that encode the flagellar structure are more-or-less expressed in the order that the products are assembled, and structural checkpoints control secretion order and timing (27,28). How and when flagella become patterned during assembly is poorly understood for any bacterium, but patterning often relies on cytoplasmic regulators similar to those that control cell division site selection (7,8).

The localization of proteins in general is thought to occur by one of two conceptual models: targeted assembly or diffusion-and-capture (29). During targeted assembly, patterning proteins determine the future localization of a complex, and the complex is assembled at that location. For instance, transmembrane proteins at one pole of the cell catalyze basal body formation in bacteria with single polar flagella (30–33). Next, the rod-cap chaperone in flagellar synthesis of Gram-negative bacteria is either fused to or associated with peptidoglycan such that rod polymerization creates holes in the peptidoglycan of sufficient diameter to allow rod transit (34–37). Finally, bushing proteins are assembled as rings around the rod to allow free rotation in the context of the envelope and govern the transition from rod-to-hook assembly (38–40). Thus, some bacteria predetermine the location of the basal body, and rod-cap activity ensures that the flagellar rod, peptidoglycan pores, and the P- and L- ring bushings are synthesized in register. During diffusion-and-capture however, proteins and/or complexes are assembled randomly, and then diffuse or otherwise move to their ultimate location where they are captured by architectural information. Whether nascent flagella are mobile prior to completion, what architectural information is recognized by flagella, and how cytoplasmic flagellar patterning proteins would govern basal body capture is unclear.

The Gram-positive bacterium *Bacillus subtilis* assembles approximately 20 flagella per cell peritrichously organized along the cell body in a non-random grid-like pattern, and appears to lack critical components of Gram-negative targeted assembly including the rod-cap, a dedicated peptidoglycan lyase, bushing proteins and transmembrane recruiters (28,40–43).

Here we explore early patterning events in *B. subtilis* by studying basal body dynamics. In wild type cells, basal bodies exhibit a negligible rate of subunit exchange, and while predominantly stationary, a low frequency of basal body mobility was observed. Using an inducible system, a high frequency of mobile basal bodies was observed soon after induction and mobility decreased over time, consistent with a model in which basal body mobility is a property of early- stage assembly and basal bodies become captured as they mature. Rod assembly is part of basal body capture as mutation of subunits early in the flagellar rod increased the frequency of mobile basal bodies, and we infer that FlgC is the rod subunit most likely to make first contact with, and be restrained by, the peptidoglycan. Finally, rod mutants also had basal body patterning defects similar to mutants defective in cytoplasmic regulatory protein FlhF that governs flagellar patterning. How *B. subtilis* flagella transit the peptidoglycan without a dedicated PG lyase to create holes sufficient to accommodate the rod and how flagella are arranged in a grid-like pattern is unclear. Our data support a model in which early stages of flagellar assembly diffuse in the membrane and metastable rod assembly continuously probes a pre-existing pattern imprinted on the peptidoglycan itself.

## RESULTS

### Flagellar basal bodies are stable and largely static

To distinguish whether flagellar patterning was primarily controlled by targeted synthesis or diffusion-and-capture in *B. subtilis*, we set out to determine if and when basal bodies were mobile. To observe flagellar basal bodies, a FliM-GFP fusion protein was incorporated into the cytoplasmic ring (C-ring) such that the flagella appear as puncta in fluorescence microscopy (41). To determine if basal bodies are mobile, cells were fluorescently imaged using time-lapse total internal reflection fluorescence (TIRF) and single-particle tracking. Approximately 5% of all basal bodies scored in the wild type were mobile over the time course observed (**Fig 1B**), and the low frequency of mobility is consistent with the fact that flagellar basal bodies have been used as stationary fiducials for comparing the movement of other proteins (44). We conclude that while the basal bodies of *B. subtilis* were overwhelmingly stationary, a low-frequency subpopulation of mobile particles exists.

**Fig 1:**
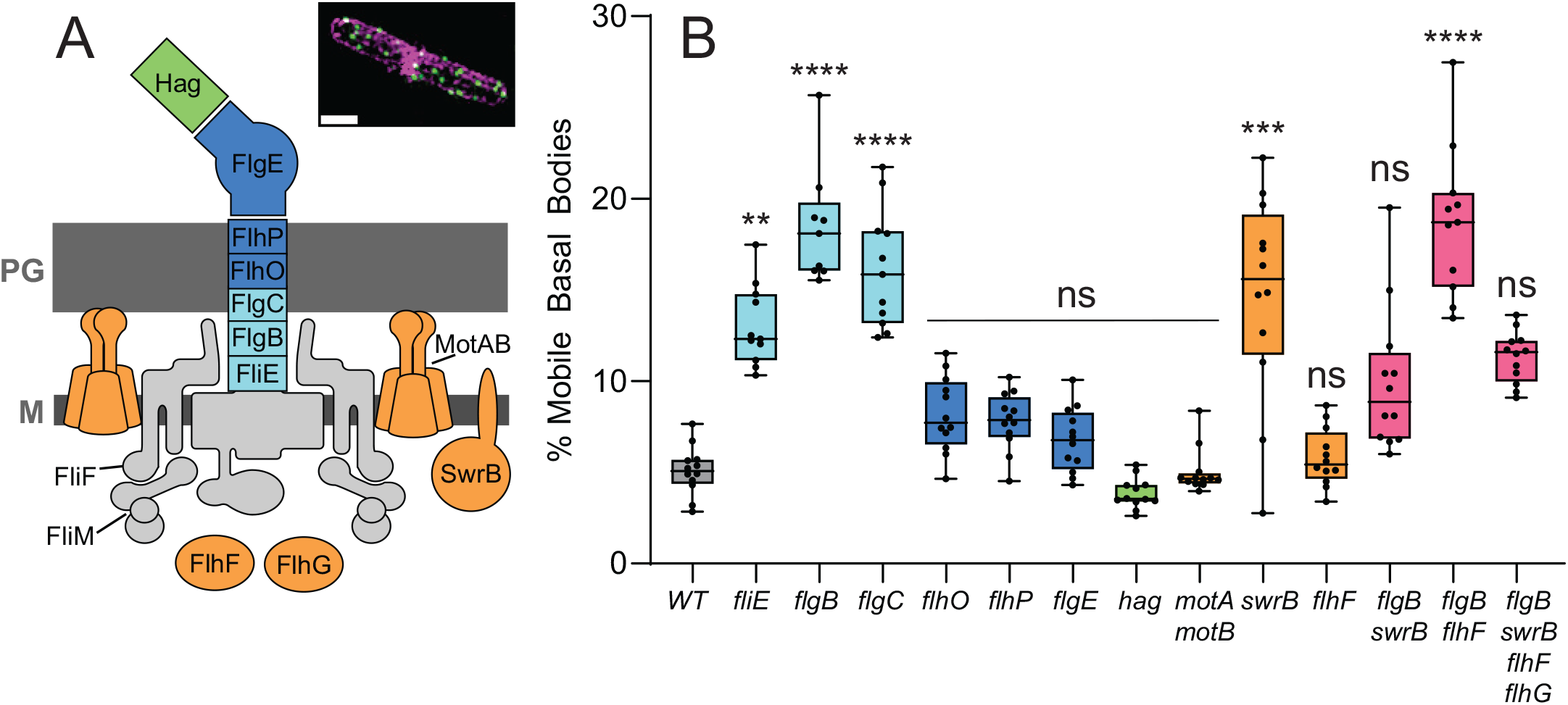
Flagellar basal bodies are immobilized by the proximal rod. (A) Cartoon cross- section of a flagellum, including the basal body and C-ring (light gray), the rod-hook (blue), filament (green), and proteins that control rotation, assembly and patterning (orange). Inset, 3D SIM fluorescence micrograph of two conjoined *B. subtilis* cells with membrane false colored red and FliM-GFP puncta false colored green. Scale bar is 1 μM. **(B)** Timelapse TIRF microscopy was used to monitor flagellar basal bodies as indicated by fluorescent puncta of FliM-GFP. Images were captured every 250 ms for 1 minute. Basal body puncta were counted, tracked, and classified as stationary or mobile based on analysis of their mean squared displacements (see methods). Three to four movies were generated for each biological replicate and scored separately to form a point on the graph. Three biological replicates were made for each strain. Boxes represent the inner quartile range of the datapoints and are color- coded according to the corresponding subunits indicated on the cartoon at the right. The line through the boxes indicates the median value and whiskers indicate the range of values measured for each dataset. The following strains were used to generate the data: WT (DK1906), *fliE* (DK5081), *flgB* (DK5082), *flgC* (DK5268), *flhO* (DK5083), *flhP* (DK5084), flgE (DK4892), *hag* (DK4347), *motAB* (DB897), *swrB* (DK479), *flhF* (DK2118), *flgB swrB* (DB1637), *flgB flhF* (DK7871), and *flgB swrB flhF flhG* (DK9675).

A low frequency of FliM-GFP puncta might indicate either that basal bodies are occasionally mobile or that C-rings might detach from the basal bodies and move independently. To test for FliM co-localization with the basal body, cells were doubly labeled in which FliM was fused to a Halo Tag and the major transmembrane flagellar housing protein FliF was fused to mNeonGreen. Although the FliM-Halo-Tag was partially functional for supporting swarming motility in single copy and the FliF-mNeonGreen fusion was non-functional (**Fig S1A**), the two fluorescent tags co-localized when expressed in the same cell and stained with a Halo reactive dye (**Fig S1B**). Moreover, rare puncta of FliM-Halo and FliF-mNeonGreen were observed to move in cells and when they did, co-localization was maintained (**Movie S1**). We conclude FliM and FliF co-localize as a complex and thus the presence of mobile puncta of FliM-GFP indicates a subpopulation of assembled and mobile basal bodies in wild type cells.

Each basal body is made up of many subunits, and even in the case of stationary basal bodies, it is possible that individual subunits are dynamically exchanged. For instance, subunits of FliM have been shown to exhibit dynamic exchange between basal bodies in *E. coli* (45–51). To determine whether individual FliM subunits were stationary or dynamic in *B. subtilis*, fluorescence redistribution after photobleaching (FRAP) experiments were performed. Briefly, puncta of FliM-GFP were bleached by exposure to localized high intensity laser excitation and fluorescence was measured at intervals by TIRF. Puncta in the laser-exposed zone of targeted cells experienced a dramatic decrease in fluorescence intensity and fluorescence intensity did not recover over time (**Fig 2A**). As a control, puncta outside the laser-exposed zone within a targeted cell experienced a minimal loss of fluorescence that was comparable to puncta in untargeted cells (**Fig 2A**). We conclude that flagellar basal bodies of wild type *B. subtilis* cells are more stable than their *E. coli* counterparts.

**Fig 2:**
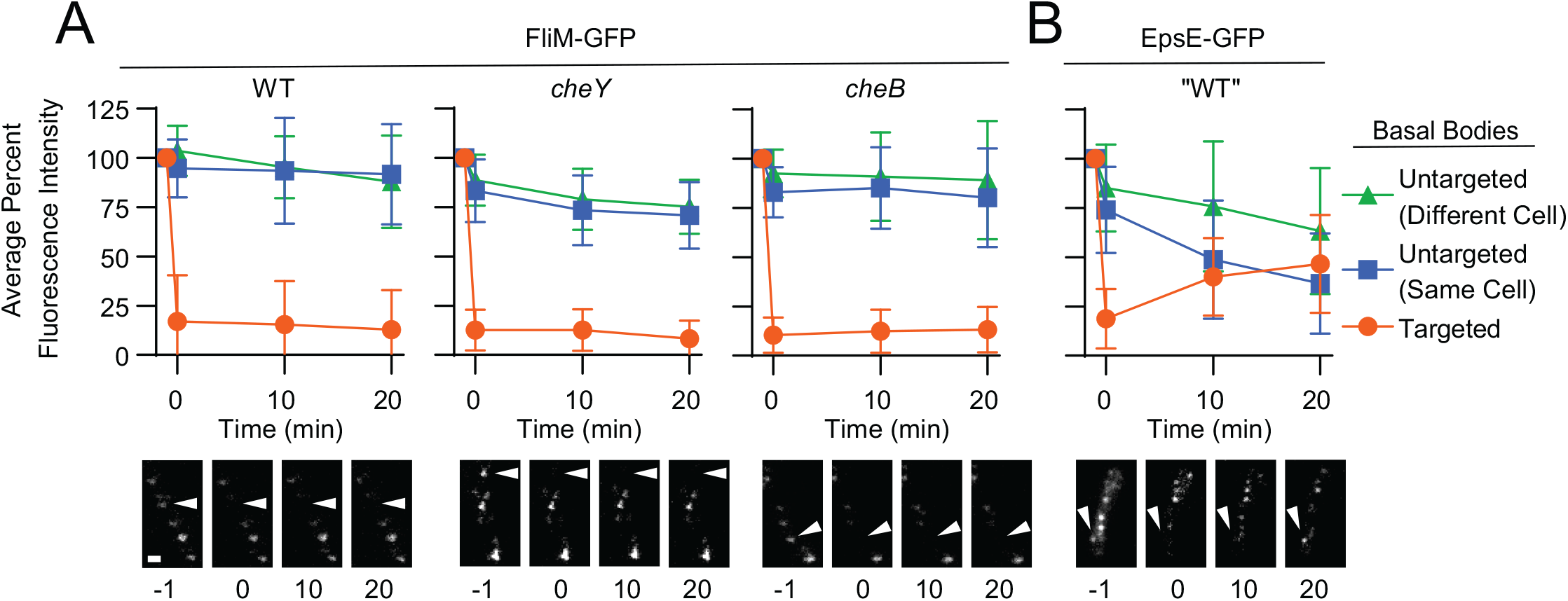
Flagellar C-ring subunits do not dynamically exchange between flagella. Cells expressing **(A)** FliM-GFP or **(B)** EpsE-GFP were imaged with TIRF microscopy before and after photobleaching in a variety of strains. After the first image was acquired, individual puncta were bleached of fluorescence via targeted laser excitation. For each strain, ten puncta were targeted for photobleaching (red circles) and fluorescence was monitored over time. In addition, fluorescence was monitored for ten puncta that were in the same cell but not targeted (blue squares) as well as ten puncta from an untargeted cell elsewhere in the same field (green triangle). Each data point represents the average % of original fluorescence intensity for 10 measurements across three biological replicates and error bars are the standard deviation. The following FliM-GFP expressing strains were used to generate panel A: wild type (DK1906), *cheY* (DK1977), and *cheB* (DK1991). For panel B, the “wild-type” strain (DS2955) was also mutated for the biofilm repressor SinR to express EpsE-GFP, and mutated for EpsH to abolish cell clumping that occurs in the absence of SinR (Kearns 2005; Blair 2008). Below each graph are representative micrographs of a single cell from each strain, with white arrows indicating the targeted basal body. Numbers indicate the time relative to laser bleaching. Scale bar is 1 μM.

The rate of dynamic exchange of FliM subunits in the *E. coli* flagellar basal body was shown to be altered by mutation of the chemotaxis signaling system (46,49,50). To determine if the seemingly stable C-rings of *B. subtilis* could be made dynamic, FRAP experiments were performed on cells in which the C-ring was conformationally locked to cause exclusive rotation in either direction by mutation of either the CheB methylesterase or the CheY the response regulator (52–55). Similar to that observed in the wild type, FliM-GFP puncta in cells mutated for CheB or CheY exhibited little to no dynamic protein exchange of individual subunits as neither fluorescence recovery of target basal bodies nor fluorescence loss of off-target basal bodies was observed after photobleaching (**Fig 2A**). Thus, it appears that although the C-rings of *E. coli* dynamically exchange structural subunits, the C-rings of *B. subtilis* do not.

While structural subunits of the *B. subtilis* flagellar C-ring appeared static, non-structural regulatory proteins have been shown to interact with the C-ring under certain conditions. For instance, EpsE is a bifunctional glycosyltransferase and flagellar clutch that inhibits flagellar rotation by binding to and disconnecting the FliG rotor component of the C-ring from power-generating stators (56–59). Unlike FliM-GFP, photobleached puncta of EpsE-GFP recovered fluorescence over time and as fluorescence recovered, off-target EpsE-GFP puncta in the same cell experienced commensurate fluorescence loss (**Fig 2B**). We conclude that regulators of C-ring function, like EpsE, can experience dynamic exchange but structural subunits of the C-ring itself do not. Thus, the majority of FliM-GFP puncta are stationary both in space and time with respect to the individual subunits that comprise the fluorescent spot.

### Flagellar basal bodies are mobile early in assembly

While the majority of basal bodies were stationary, a minority were mobile. We hypothesized that the mobile subpopulation represented immature basal bodies at an early stage of assembly. To observe the behavior of immature basal bodies, a strain was created in which basal body formation could be controlled by placing the *fla/che* operon under the control of an IPTG-inducible promoter (Guttenplan 2013). Thus in the absence of inducer, the strain should contain no flagellar basal bodies and all basal bodies observed after inducer addition would be newly synthesized (**Fig 3A**). In this construct, the native *fliM* gene was deleted and a FliM-GFP construct was expressed from the *P_fla/che_* promoter at an ectopic site (41) such that FliM-GFP would be present prior to IPTG-induction and puncta formation would be restricted by nucleation rather than expression (**Fig 3A**). Cells were monitored by two different approaches. In one approach, the total number of basal bodies was counted at indicated time points using OMX 3D SIM that resolves fluorescence originating in many planes of the cell. In a complementary approach, time-lapse TIRF microscopy was used to focus on a single plane closest to the coverslip to isolate and quantify basal body mobility by single-particle tracking. Combined, the two approaches give both the number of basal bodies per cell plus the proportion of basal bodies that were mobile.

**Fig 3:**
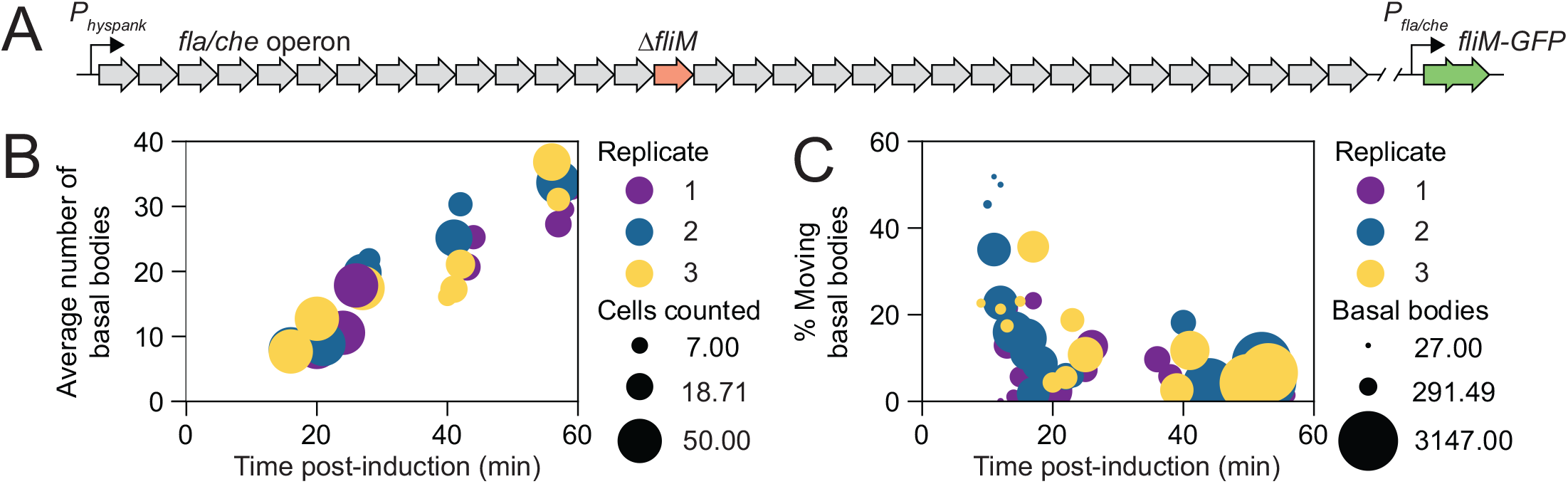
Flagellar basal bodies are mobile early in assembly. (A) Schematic showing the *fla/che* operon with the IPTG-inducible *P_hyspank_*promoter replacing the native *P_flache_* promoter. In addition, the *fliM* gene was deleted from the *fla/che* operon and ectopically reintroduced as *fliM- GFP* under the control *P_flache_*. **(B)** Cells described in panel A (DK31) were imaged with 3D SIM at various timepoints after the addition of IPTG. The number of FliM-GFP puncta were counted to determine the number of basal bodies for fifty cells in each of three biological replicates, totaling 150 cells. Each bubble on the plot shows the average number of basal bodies per cell in one individual image with the size of the bubble representing the number of cells in that image. **(C)** The same strain was imaged with timelapse TIRF microscopy at varying timepoints after the addition of IPTG. Images were captured every 250 ms for 1 minute. Basal body puncta were counted, tracked, and classified as stationary or mobile based on analysis of their mean squared displacements. Three to four movies were generated for each biological replicate and scored separately to form a bubble on the plot. The size of the bubble represents the number of basal bodies identified in that movie.

In the absence of inducer, cells containing the IPTG-inducible *fla/che* operon grew as long chains and FliM-GFP signal was diffuse in the cytoplasm consistent with a failure to express the *fla/che* operon and consequently a failure to produce nascent basal bodies upon which FliM-GFP would nucleate (41,60–62). Next, IPTG was added at mid-log phase and cells were observed at various timepoints by OMX-3D SIM and TIRF microscopy in separate experiments. Basal body puncta were observed at the earliest timepoints tested, and consistent with temporal assembly, the number of puncta increased over time (**Fig 3B**). Meanwhile, the frequency of basal body mobility was highest at the earliest timepoints, and decreased over time (**Fig 3C**). We conclude that newly synthesized basal bodies exhibit the greatest mobility and that the frequency of mobility decreases as the basal bodies mature. We infer that at least part of flagellar patterning involves movement of immature flagellar basal bodies that are captured at a location later in flagellar assembly.

### Flagellar basal bodies are immobilized by synthesis of the flagellar rod

To determine the mechanism by which flagellar basal bodies are captured and immobilized, FliM-GFP puncta dynamics were measured by TIRF timelapse microscopy in a variety of mutants. The stage in flagellar assembly that follows the formation of the basal body is the secretion and polymerization of subunits that make up the flagellar rod and hook (12,13,19,23,63,64). The flagellar rod-hook is comprised of six different proteins which are sequentially assembled from most proximal to distal in the following order: FliE, FlgB, FlgC, FlhO, FlhP and FlgE (24,65,66) (**Fig 1A**). Mutation of FliE and the proximal rod subunits FlgB, and FlgC increased the frequency of basal body mobility significantly over wild type (**Fig 1B**). Mutation of the distal rod subunits FlhO, and FlhP, and the hook subunit, FlgE, appeared to increase basal body mobility but the increase was not significantly different from wild type, perhaps because significance was determined by a Kruskal-Wallis test. We note that Kruskal-Wallis is less sensitive/powerful than a more standard ANOVA test, but the ANOVA test is inappropriate when comparing data expressed as percentages (67). The flagellar rod-hook of *B. subtilis* is metastable in that it both assembles and disassembles until the transition to the flagellar filament (66), and this property may account for the apparent, but not statistically significant increase, in basal body mobility in some mutants. Finally, mutation of the flagellar filament subunit protein Hag, assembled after completion of the rod-hook, did not increase basal body mobility above wild type levels (**Fig 1B**). We conclude that mutation of the proximal rod increases the frequency of basal body mobility to approximately 20% at steady state.

While rod transit through peptidoglycan appears to contribute substantially to immobilization of the basal body, we note that as many as 80% of the basal bodies disrupted for the proximal rod appeared to remain stationary (**Fig 1B**). To explore whether non-structural regulatory proteins also contributed to basal body mobility, we focused on regulators involved in flagellar function and assembly. MotA and MotB form the stator complex that binds to peptidoglycan and revolves in interaction with the C-ring to power flagellar rotation (68–72).

Mutants simultaneously disrupted for MotA and MotB did not increase basal body mobility above wild type levels, consistent with the fact that the stators are a late-acting component and likely associate after flagella have been fully assembled and patterned (**Fig 1B**). SwrB is a single-pass transmembrane protein with a cytoplasmic domain that promotes basal body maturation and activation of the flagellar type III secretion system (23,73). Mutation of SwrB significantly increased basal body mobility above that of wild type resembling cells mutated for the early rod subunits, and simultaneous mutation of SwrB and the early rod subunit FlgB did not further increase basal body mobility (**Fig 1B**). We conclude that SwrB likely immobilizes basal bodies by activating flagellar type III secretion and promoting proximal rod subunit export.

A third regulator we tested was FlhF and FlhG, partner proteins that control flagellar patterning. FlhG is a MinD-like ATPase which when mutated, abolishes distance control between basal bodies and causes basal body aggregation in the cell (41,74–76). While mobile basal bodies were occasionally observed in *flhG* mutant cells, the frequency of basal body mobility could not be determined as basal body aggregation confounded puncta counting (**Movie S2**). FlhF is an SRP-like GTPase thought to interact with FlhG, and mutation of FlhF causes an asymmetric flagellar distribution that skews towards one cell pole (41,74,77,78). A mutant disrupted for FlhF did not significantly increase basal body mobility above wild type levels, and simultaneous mutation of FlhF and FlgB, or FlgB, SwrB, FlhF, and FlhG did not increase basal body mobility above mutation of FlgB alone (**Fig 1B**). We conclude that FlhF and FlhG do not control patterning by regulating the frequency of mobile basal bodies. Thus, the flagellar rod appears to be the primary determinant of basal body immobilization.

### The flagellar rod is required for symmetrical patterning

Mutations that disrupted the proximal flagellar rod increased basal body mobility but mutations that disrupted the patterning regulators FlhF and FlhG did not. To further explore the relationship between mobility and patterning, each strain was imaged using OMX-3D SIM (**Fig 4A**) and the nearest neighbor mean distance (NNMD) was determined and used to calculate a Clark-Evans ratio (41) (**Fig 4B**). A Clark-Evans ratio varies between 0 and 2 where a value of 1 indicates a random distribution and values near 0 or 2 indicates a non-random distribution with either clumped or grid-like quality, respectively (79). In wild type, each basal body is synthesized with an NNMD greater than that which would be predicted by chance (41), and the Clark-Evans ratio was between a value of 1 and 2, consistent with a grid-like distribution (**Fig 4B**). Each mutant tested in this study exhibited an NNMD similar to that of the wild type and the Clark-Evans ratios were also of values greater than 1 (**Fig 4, Fig S2**). Thus, whether or not a mutation increased basal body mobility, the overall pattern of flagellar placement appeared to be largely conserved, at least at the level of the distance between basal bodies.

**Fig 4:**
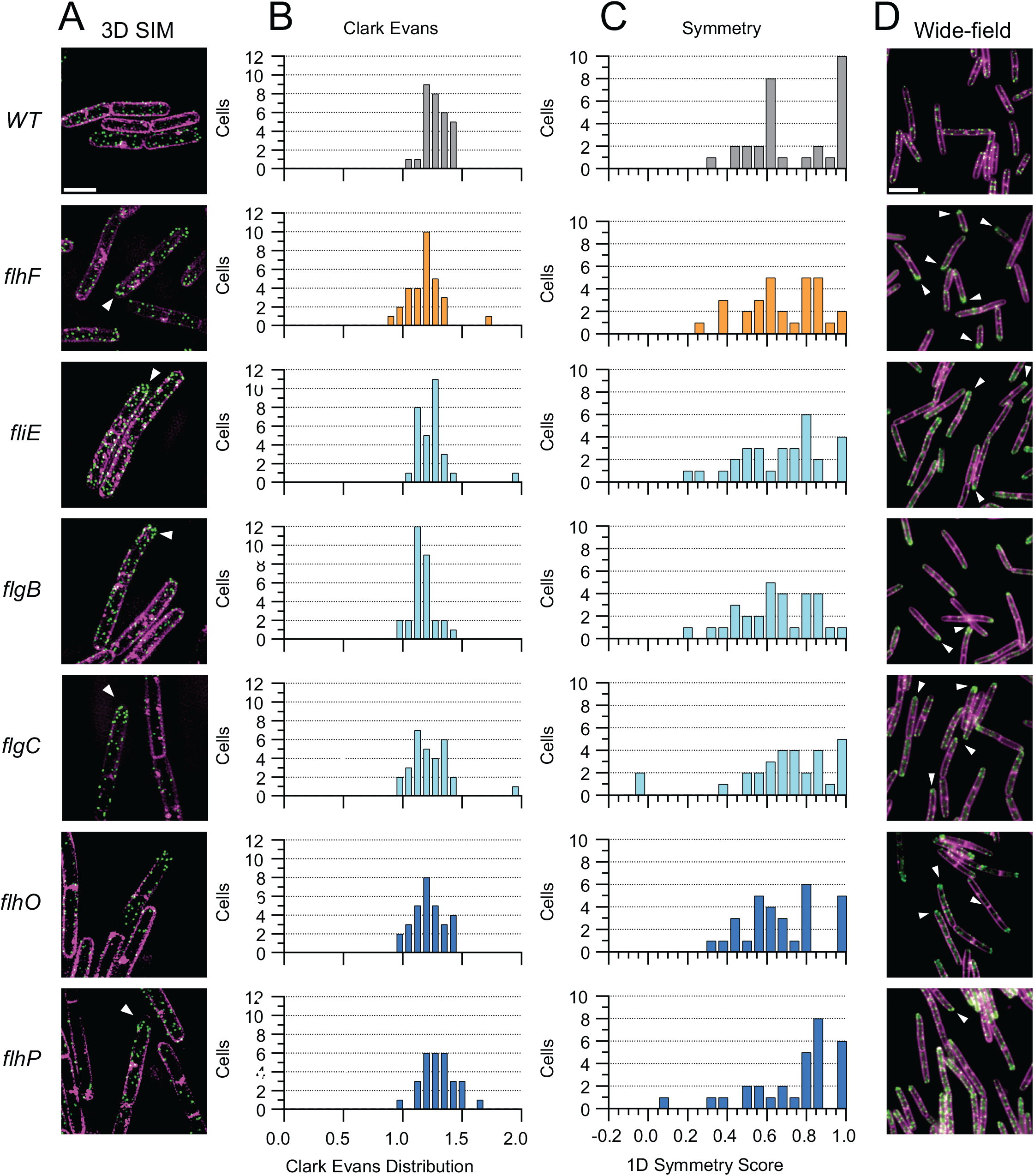
The flagellar rod is required for basal body patterning. (A) 3D SIM images of FliM-GFP (green) representing the flagellar basal body and FM4-64 (magenta) showing the cell membrane. Scale bar is 2 μM. **(B)** Clark-Evans distributions were calculated from the NNMD score from each cell and projected as a histogram of individuals to evaluate patterning of flagellar basal bodies (see methods). The data from 30 cells are included in each graph, in which 10 cells were counted in each of three replicates. **(C)** To measure the symmetry of basal bodies in each cell, the number of basal bodies were counted in each half of the cell, the smaller number was divided by the larger number, and the data was presented as a histogram. Thus, a value of 1 indicates a perfectly symmetrical distribution, while values less than one indicate asymmetry. The data from 30 cells are included in each graph, in which 10 cells were counted in each of three replicates. **(D)** Wide field images of FliM-GFP (green) representing the flagellar basal body and FM4-64 (magenta) showing the cell membrane. White carats indicate polar accumulation of FliM-GFP. Scale bar is 5 μM. The following strains were used to generate the data: WT (DK1906), *flhF* (DK2118), *fliE* (DK5081), *flgB* (DK5082), *flgC* (DK5268), *flhO* (DK5083), and *flhP* (DK5084).

**Fig 5:**
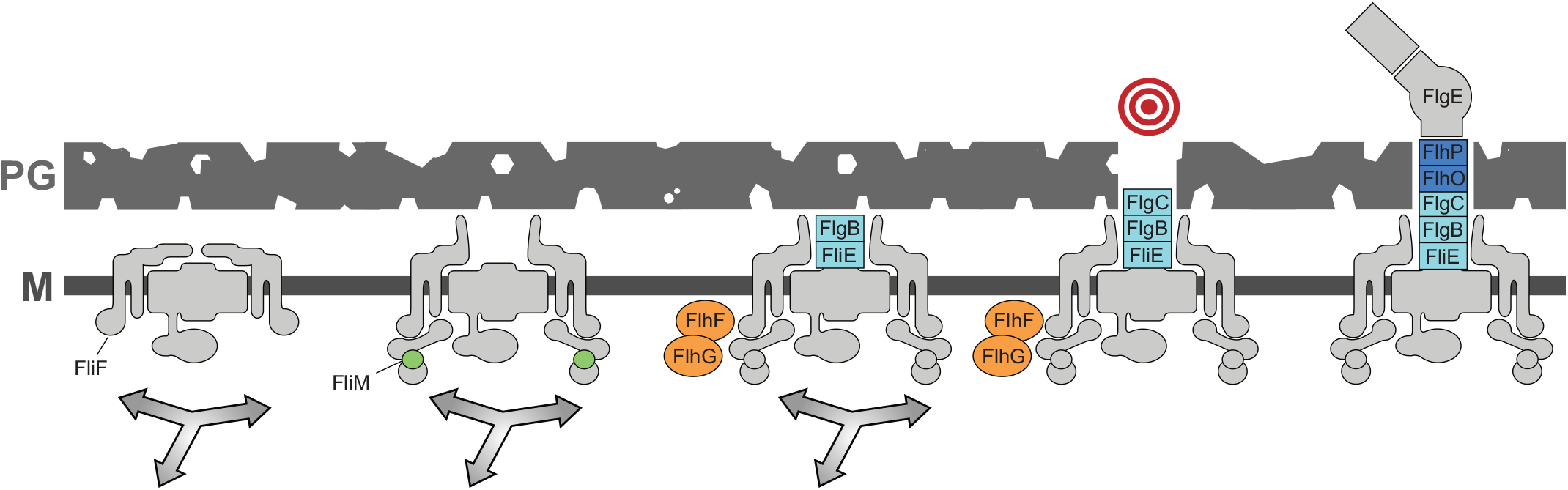
Basal bodies are immobilized by rod assembly. A model of the steps in dynamic flagellar assembly. Basal bodies are mobile (arrows) but inactive, and become active for secretion by assembly of the C-ring (FliM, colored green). Next, rod subunits are secreted and polymerized on top of the basal body to form the proximal rod. If the local peptidoglycan (PG) superstructure does not sterically permit assembly of the full rod, rod subunits disassemble to a point below the steric hindrance of the PG, and remobilize the basal body. Thus, the rod probes the PG for permissive holes and if a hole is found of sufficient diameter, rod-hook assembly will be completed, the basal body will transition to filament subunit export and assembly, and lock the entire structure in place. The patterning proteins FlhF and FlhG (colored orange) interact with FliF and the C-ring respectively and either expedite basal body search or restrict which holes may be occupied, to create a symmetrically-distributed, grid-like pattern of flagella.

We noted that while the NNMD measurements did not indicate a dramatic difference between basal body patterning in the mutants (**Fig S2**), many of the mutants had a Clark-Evans ratio that appeared to skew closer to a value of 1 indicating a more random distribution in some of the individuals (**Fig 4B**). One such mutant was the strain defective for the patterning protein FlhF which had previously been shown to display an asymmetric distribution of basal bodies and frequent accumulation of basal bodies towards one cell pole (41) (**Fig 4B**). To measure symmetrical distribution, a 1D symmetry score was calculated by splitting cells imaged by OMX 3D-SIM in half, counting the number of basal bodies per side and dividing the lower value by the higher. Wild type cells had a high degree of symmetry in basal body distribution (values close to 1) but the *flhF* mutant exhibited less symmetry, consistent with previous results (**Fig 4C**) (41). The symmetry values of each of the rod mutants resembled that of cells defective for FlhF, again with a broader and flatter distribution and an overall reduction in the number of cells with a value of 1 (**Fig 4C**). Finally, many cells in the populations of strains defective in FlhF and the rod subunits also appeared to accumulate basal bodies at one pole, a phenomenon more easily observed in conventional fluorescence microscopy (**Fig 4D**). We conclude that mutation of the extracytoplasmic flagellar rod subunits confers defects in patterning that are similar to defects observed in the cytoplasmic FlhF flagellar patterning protein. We further conclude that the rod both immobilizes basal bodies and participates in pattern formation.

## DISCUSSION

How flagella are positioned on the bacterial cell surface is poorly understood, and here we explore the earliest stages of pattern acquisition in *B. subtilis*. We show that basal bodies are predominantly stationary at steady state but are mobile early in their synthesis using fluorescently labeled FliM as a proxy. As FliM loading and C-ring completion is required for basal body maturation and activation of the secretion system (21–23,80,81), puncta of FliM allow observation of events between the stages of basal body and rod assembly. Mobile basal bodies become immobilized and patterned as they mature, and both patterning and immobilization depend on structural subunits of the flagellar rod as they transit the peptidoglycan. While the average pore size in peptidoglycan is smaller than the diameter of the rod (21,64,82,83), new observations and models of peptidoglycan superstructure invoke heterogeneous porosity (84–87). We argue that basal bodies diffuse in the membrane until rod polymerization is permitted at a pore of sufficient diameter thereby capturing the complex. Thus, flagella of *B. subtilis* search for holes in the wall rather than making them, and cytoplasmic patterning systems somehow restrict which holes may be occupied.

Our model incorporates the features of the flagellar rod that differentiate it from the hook and filament. First, the hook and the filament are made of single proteins FlgE (66,88,89) and Hag (FliC) (90–92) respectively, but the rod is comprised of the basal body connector FliE (93,94) and four different homologous structural subunits FlgB, FlgC, FlhO (FlgF), and FlhP (FlgG) (11,66,95). Rod subunits are secreted through the rod itself and sequentially stacked in helices of 6-24 subunits (11,97), and our data suggest that FlgC, as the most distal subunit necessary to immobilize the basal body, is likely the first subunit to contact peptidoglycan.

Second, the hook and filament are polymerized by dedicated cap chaperones (FlgD and FliD, respectively) (98,99), and Gram-negative bacteria encode a dedicated rod cap/PG lyase complex (FlgJ) thought to make holes for rod transit (34–36,100). *B. subtilis* however, seems to lack both a rod cap and a dedicated flagellar PG lyase, rendering the model in which the rod creates holes in the PG seemingly inapplicable (28,42,43). Third, whereas the hook and filament are stable structures, the rod is metastable, meaning it can both polymerize and depolymerize (a phenomenon observed primarily when completion is impaired) (12,13,19,66). We speculate that if the rod is prevented from completion due to steric constraints imposed by the PG, rod assembly will stall, depolymerize until the basal body is remobilized, and relocate for another attempt. In short, we suggest that stochastic rod assembly probes the peptidoglycan superstructure. The flagellar secretion system exports approximately 10 rod subunits per second, so sampling and relocating could be rapid (101,102).

While flagellar rod assembly immobilizes approximately 20% of basal bodies, ∼80% remained immobile after rod ablation. Some basal bodies may appear immobile for technical reasons. Perhaps all basal body puncta are mobilized in a rod mutant but some move too slowly to detect displacement within our experimental parameters, and we note that basal bodies in *E. coli* have been reported to move in a manner consistent with both free and restricted diffusion (103). We further note that membranes are heterogeneous and basal bodies are large multi-subunit complexes that could be stochastically distributed between regions of high and low fluidity (104–106). Furthermore, experimental parameters such as agarose pad firmness or composition may affect mobility, and we found that mobility decreased with intense fluorescence excitation as in the case where multiple fluorophores were simultaneously imaged over long time periods. Alternatively, rod-independent immobilization might be mediated by FliF, part of the flagellar secretion system, or an as-yet-undiscovered protein. Whatever the case, the predominance of basal body immobility permitted pattern analysis, even when partially mobilized by rod mutation.

The connection between flagellar mobility and patterning is most clearly observed when flagella synthesis is artificially induced. Using an inducible system, basal bodies were mobile at high frequency soon after induction, and the frequency of mobility decreased with time, consistent with a diffusion-and-capture model for flagellar patterning. After 40 minutes of induction, both the number of basal bodies and pattern is similar to that of the wild type at steady state (**Fig 3B**, **Fig S3**). Thus we infer that whatever restricts basal body mobility and patterning is either encoded within the induced flagellar regulon itself and rapidly imprinted, or is exogenous to the flagellar regulon and pre-existent at the time of induction. The two possibilities are unified in our model in which rod protein assembly searches a pre-existent pattern of holes intrinsic to the peptidoglycan and flagellar rod mutants not only increase basal body mobility but display a patterning defect. The rod alone is insufficient for patterning, however, as the cytoplasmic proteins FlhF and FlhG disrupt pattern formation but synthesize flagella and are motile.

FlhF and FlhG are cytosolic proteins that govern flagellar patterning in a wide variety of bacterial species. FlhF is thought to interact with the basal body protein FliF and in *B. subtilis*, FlhF inhibits polar accumulation of flagella (41,107,108). FlhG is thought to interact with the C-ring proteins FliM and FliY and in *B. subtilis* FlhG controls basal body NNMD to inhibit basal body aggregation (41,75,109). Here we show that mutation of the flagellar rod phenocopies mutation of FlhF for asymmetric flagellar distribution. Why mutation of intracellular regulators and extracellular structural subunits phenocopy for patterning defects is unclear but supports a sensory/regulatory role for rod assembly. We infer that the assembly state of the rod could be transmitted to the cytoplasm through conformational changes in the basal body with which FlhF and FlhG interact. How FlhF/FlhG restrict rod formation to a subset of permissible holes in the peptidoglycan to create the grid-like symmetrical distribution of mature flagella remains unknown.

## MATERIALS AND METHODS

### Strains and growth conditions

*B. subtilis* strains were grown in lysogeny broth (LB) (10 g tryptone, 5 g yeast extract, 5 g NaCl per L) broth or on LB plates fortified with 1.5% Bacto agar at 37°C. When appropriate, antibiotics were included at the following concentrations: 10 µg/ml tetracycline, 100 µg/ml spectinomycin, 5 µg/ml chloramphenicol, 5 µg/ml kanamycin, and 1 µg/ml erythromycin plus 25 µg/ml lincomycin (*mls*). Isopropyl β-D-thiogalactopyranoside (IPTG, Sigma) was added to the medium at the indicated concentration when appropriate.

#### Strain construction

All PCR products were amplified from *B. subtilis* chromosomal DNA, from the indicated strains. All constructs were either transformed into a NCIB3610-derived natural competent strain DK1042 (110), or transformed into domesticated strain PY79 and then moved to the 3610 background using SPP1-mediated generalized phage transduction (111). Briefly, SPP1-mediated transduction was performed by generating a plaque lysate on *B. subtilis* grown in TY soft agar (1% tryptone, 0.5% yeast extract, 0.5% NaCl, 10 mM MgSO_4_, and 1 mM MnSO4, 0.5% Bacto agar). Recipient strains were grown to stationary phase in TY, 1 ml was diluted into 9 ml TY, and 25 µl lysate was added, followed by incubation at room temperature for 30 min and then selection on the respective antibiotic at 37°C overnight. For transductions in which spectinomycin, kanamycin, or chloramphenicol resistance was selected for, 10 mM sodium citrate was added to the selection plate. All strains used in this study are listed in Table 1. All primers used in this study are listed in Table S2.

**Table 1:**
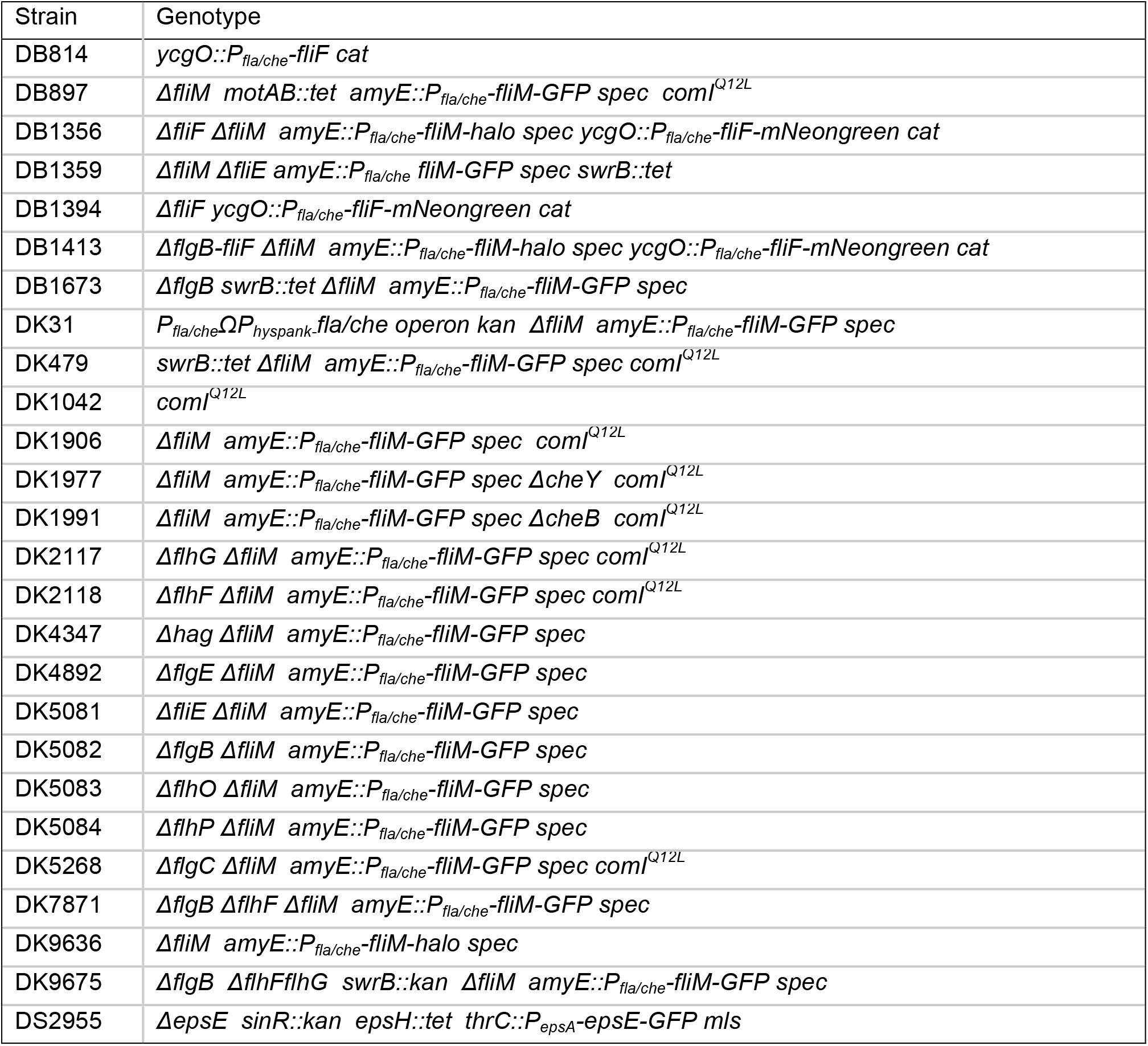
Strains.

To generate the translational fusion of FliF to mNeonGreen, the region containing the 5’ end of *ycgO*, the *P_flache_* promoter, and *fliF* was amplified from strain DB814 using primer pair 7620/8303, and the region containing the mNeongreen gene, the cat gene conferring chloramphenicol resistance and the 3’ end of *ycgO* was amplified from strain DB1341 using primer pair 8307/7621. The two fragments were fused by Gibson assembly and transformed into *B. subtilis*.

To generate pCD9 in which *fliM* was expressed from the *P_flache_* promoter with a convenient site for cloning 3’ end fusions, the *P_fla/che_-fliM* construct was amplified from pSG49 (41) using primer pair 8518/8519, digested with SphI and NotI and cloned into the SphI-NotI sites of pAH25 encoding a spectinomycin resistance cassette between two arms of the amyE gene (a generous gift from Amy Camp, Mount Holyoke College),.

To generate pDP580 which encodes a translational fusion of FliM to halo, the gene encoding halo was PCR amplified from an amplicon (generous gift of Malcolm Winkler, Indiana University) using primers 7681/7682, digested with NheI and NotI and cloned into the NheI-NotI sites of pCD9.

### Swarm expansion assay

For the swarm expansion assay, cells were grown to mid-log phase at 37°C in LB broth, pelleted, and resuspended to 10 OD_600_ in pH 8.0 phosphate-buffered saline (PBS) (137mM NaCl, 2.7mM KCl, 10mM Na_2_HPO_4_, and 2mM KH_2_PO_4_) containing 0.5% India ink. Freshly prepared LB containing 0.7% Bacto agar (25 ml/plate) was dried for 10 min in a laminar flow hood, centrally inoculated with 10 μl of the cell suspension, dried for another 10 min, and incubated at 37°C. The India ink demarks the origin of the colony and the swarm radius was measured relative to the origin. For consistency, an axis was drawn on the back of the plate and swarm radii measurements were taken along this transect. Swarm expansion was measured every 30 minutes until wild-type reached a radius of 30mm (54).

### Preparation of cells for microscopy

Cells were grown in LB broth at 37C until log phase (OD_600_ 0.5-0.9). For halo-tag microscopy, cells were incubated in LB broth at 37C with 1 nM Janelia Fluor 549 for 10 minutes. One milliliter of cells was then pelleted. For 3D SIM microscopy, cells were resuspended in 30 μL 5 μg/ml FM4-64 (Molecular Probes) and incubated at room temperature for 3 minutes to stain the membrane before being pelleted again. Cells were then washed with 1 mL water and pelleted again. Samples were resuspended in 30 μL water, then observed by spotting 4 μL of this suspension on a 1% agarose pad made with water. For FRAP microscopy, water was replaced by the defined rich casein hydrolysate (CH) media in all instances. (112)

### OMX microscopy

A GE Deltavision OMX 3D-SIM Super Resolution System V3.0 (Applied Precision) was used for Fluorescent Redistribution After Photobleaching (FRAP) experiments, Total Internal Reflection Fluorescence (TIRF) microscopy, and 3D Structured Illumination Microscopy (SIM). Images were acquired with the PCO.edge front illuminated sCMOS camera (PCO-Tech, Wilmington Delaware). Image acquisition was directed by AcquireSR 4.5 (Applied Precision, Cytiva). For TIRF and FRAP, a 1.49NA Apo N 60X oil objective was used with 1.514 refractive index oil. For 3D SIM, a 1.42NA PlanApo 60X oil objective was used with 1.520 refractive index oil. Two channel images were acquired sequentially using the 568 (609–654 emission filter) and 488 (500–550 emission filter) nm lasers (Toptica, Model: iCHROME-MLE-LFK-GE). Single channel images were captured using the 488 nm laser alone. Exposure times were 15-30ms. 25-30% of the original 100mW (488nm) or 65mW (568nm) laser power was used for image acquisition. For Timelapse TIRF, images were taken every 250 milliseconds for one minute. Supplemental movies are played back at 60 frames per second. For 3D SIM, image reconstructions were made with the OMX specific softWoRx 7.2.2 suite (DeltaVision).

### Basal body distribution analysis from 3D SIM images

Basal body position was determined using particle identification tools in Imaris Bitplane software on 3D SIM files from the OMX microscope. The X, Y, and Z positions were determined by model fitting to a sub-diffraction object after entering information about the imaging conditions and microscope parameters. The X and Y positions of individual cells were selected by hand using membrane staining as fiducial reference for cell axes. Basal body positions were exported in calibrated micrometers to Excel spreadsheets.

Basal body positions were imported into MATLAB and associated into matrices for X, Y, and Z positions and for cell ends. Estimates of basal body distribution on the bacterial cell perimeter were created using the assumption that cells are roughly cylindrical. Each cell was centered by subtracting the mean X, Y, and Z basal body position for each respective dimension. Cells were rotated to have the long axis on X by creating a definite positive 3x3 matrix from the centered X/Y/Z positions and using singular value decomposition to create a rotation matrix. Because basal body positions were initially used to center the cell, the end-wall positions were applied to recenter the cell’s long axis to the cell’s midpoint. The basal body asymmetry over the long axis was determined as the ratio of higher to lower basal body count relative to the cell midpoint.

Mean basal body distribution was determined by normalizing cell lengths and creating a histogram of 21 equal bins for relative basal body position with cells randomized for orientation (i.e., no preference for basal body asymmetry).

Basal body displacement was determined essentially as in (prior cite). A cylinder was created by fitting a circle to Y/Z projected basal body positions and then extending the 3D cylinder over the X axis. Nearest neighbor distance was calculated using the projected position of each basal body on the ‘unrolled’ cylinder. Spatial distribution bias was estimated using a Clark Evans formulation (cite) with the cylinder surface (exclusive of cell ends) used for area estimation when calculating 2D density. The spatial bias estimation is a ratio of mean nearest neighbor distance and an assumed random (Poisson) 2D distribution over the same surface area yielding a range from 0 (clumped) to 2 (maximally dispersed) with 1 representing a random distribution (cite). Cell volume was calculated as volume of the cylinder.

### Fluorescent redistribution experiments

For Fluorescence Redistribution after Photobleaching (FRAP) experiments, cells were imaged immediately before and after the bleaching event, and for the ten- and twenty- minute timepoints using the TIRF settings described in the “OMX Microscopy” methods section. For the bleaching event, regions of cells were exposed to a 488 nm laser moving in a Lissajous pattern for a duration of 0.4 seconds. Cells in which some, but not all or most, of their foci bleached, were analyzed for fluorescence redistribution.

For each biological replicate within each strain, ten regions of interest (ROIs) were drawn randomly on the field background, where no cells were present. Fluorescence intensity was measured at each timepoint (immediately before and after bleaching, then at ten- and twenty- minutes post bleaching) and these were averaged to give a measure of background at each timepoint. Next, ROIs were drawn around ten bleached puncta, ten unbleached puncta in the same cells as the previous ten, and ten unbleached puncta in cells with no localized bleaching. The fluorescence intensity of these spots were measured at each timepoint. The average background at a given timepoint was subtracted from the fluorescence value of each measured punctum. These values were then averaged. The original average fluorescence intensity was set as 100% and the following values were reported as a percentage of the original averaged value.

### Basal body tracking and mobility quantification

Movies of basal bodies with fluorescently labeled FliM-GFP subunits were analyzed with Palmari, a particle tracking plugin for Napari (113). Prior to analysis, movies were reviewed and those with substantial drift or cell movement that would affect particle tracking were excluded. At least three movies were considered per strain or condition. A mask determined by an intensity threshold on the time average of the entire movie was used to exclude regions without cells from analysis. Particles were detected by applying a Laplacian of Gaussian filter to each frame and identifying particles with intensities greater than one standard deviation above the mean intensity for that frame (114). For *fla/che* induction experiments, the intensity threshold was adjusted to account for the wide variance in particle densities. The centroid position of each particle was estimated using radial symmetry optimization (115). Particle localizations were linked into tracks using a simple tracking algorithm that links particles detected in consecutive frames within a 3-pixel search radius (116).

To quantify the mobility of labeled basal bodies, we considered tracks with localizations in at least 20 consecutive frames. Tracks were analyzed in 20-frame segments on a rolling basis. Mean squared displacements, 〈𝑟^2^〉, for all temporal lags, 𝜏, within a segment were used to estimate the apparent diffusion coefficient, 𝐷_𝑎𝑝𝑝_. For Brownian motion, 〈𝑟^2^〉(𝜏) is proportional to 𝐷_𝑎𝑝𝑝_:

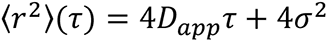

where 𝜏 is an integer multiple, 𝑖, of the 250-ms time interval between frames. 𝐷_𝑎𝑝𝑝_ was estimated using an analytical expression from weighted least squares optimization (117):

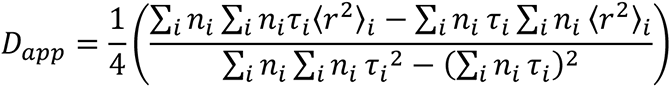

Here, 𝑛_𝑖_ is the number of displacements used to calculate the mean squared displacement, 〈𝑟^2^〉_𝑖_, for a temporal lag, 𝜏_𝑖_.

We determined that 0.0003 µm^2^/s was the lower limit of detectable 𝐷_𝑎𝑝𝑝_ based on the localization uncertainty of these experiments. Particles that were considered mobile if their apparent diffusion coefficient exceeded 0.0003 µm^2^/s in at least 10% of segments.

Analysis was completed using a series of IPython notebooks that have been made available in a Github repository: https://github.com/d-foust/basal_body_tracking.

## ACKNOWLEDGMENTS

We thank Jim Powers and Andras Kun assistance at the Indiana University Light Microscopy Imaging Center (LMIC). Support for this work comes from National Institutes of Health grants R35-GM131783 to DBK, R01-GM144731 to JSB, NIH1S10OD024988-01 to the Indiana

University Light Microscopy Imaging Center (LMIC), and National Science Foundation grant MCB1927504 to SS.

